# Now you see it: fixation-related electrical potentials during a free visual search task reveal the timing of visual awareness

**DOI:** 10.1101/2022.11.02.514808

**Authors:** Zeguo Qiu, Hongfeng Xia, Stefanie I. Becker, Zachary Hamblin-Frohman, Alan J. Pegna

**Author notes:** **Corresponding author** Alan J. Pegna, School of Psychology, The University of Queensland, Brisbane 4072, Australia.

## Abstract

It has been repeatedly claimed that emotional faces capture attention readily, and that they are processed without awareness. Yet some observations cast doubt on these assertions. Part of the problem may lie in the experimental paradigms employed. Here, we used a free viewing visual search task and simultaneously recorded electroencephalography and eye-movements. Fixation-related potentials were computed for fearful and neutral facial expressions, and the electrical response compared when participants were aware or unaware of the fixated stimulus. We showed that the P300 increased across repeated fixations on the unseen targets, culminating in a conscious report, likely reflecting evidence accumulation. Awareness of the stimulus was associated with electrical changes emerging at around 130 ms, with emotions of the stimulus being dissociated only after awareness had arisen. These results suggest that the earliest electrical correlate of awareness emerges at around 130 ms in visual search and that emotion processing requires visual awareness.

## Main

Although the visual field of humans spans nearly 180°^1,2^, effective visual processing occurs only at the centre, in the so-called functional visual field^3^. Consequently, when exploring their visual environment, humans will produce multiple saccades and fixations, orienting their gaze towards different parts of the visual field to process the stimuli that capture their attention or are relevant to their goals^4,5^. In such visual search tasks, it has long been observed that the exploration time increases as a function of the number of items in the visual field^6^. Variations in the speed of detection have been found across stimuli, which are shown to depend on a number of parameters including the physical characteristics of the stimuli and the features that distinguish the stimuli from irrelevant distractors^7^.

While central fixation is important for conscious processing, it is not sufficient to generate awareness. Indeed, when exploring the environment, stimuli may be fixated without being consciously detected, and decades of behavioural experiments using visual search paradigms have shown that, during serial search, targets may be fixated without being consciously seen^8^.

Human faces, in particular those displaying emotional expressions, constitute a particularly important category of stimuli in that they are crucial to our daily social life. Emotional faces have received much scrutiny following early observations indicating that they can readily capture attention^9,10,11^. Behavioural studies examining the speed of visual search and detection have reported that emotional faces are detected faster than other types of stimuli^12,13,14^. In line with these observations, studies of patients with spatial attention deficits have shown that emotional faces attract attention more efficiently than non-emotional stimuli^11,15^, possibly due to their behavioural relevance. In a similar vein, patients with visual deficits have been found to process emotional faces without awareness (a phenomenon termed affective blindsight^16,17^). This finding has been replicated in healthy controls using visual masking^16,17,18,19,20^.

However, objections have been voiced suggesting that the above-mentioned attentional effects may be driven essentially by the low-level characteristics of the stimuli^21^, while others have argued that they occur only if sufficient attentional resources are available, and that increasing the attentional requirements of a concurrent task prevents faces from capturing attention^22^. Unconscious viewing has also been questioned and suggested to be a degraded form of conscious vision^23,24^.

Recently, neuroscience has begun to explore in more depth the neural events that reflect perceptual awareness, as well as their temporal dynamics (see Mudrik and Deouell^25^ for a recent review). Using electrophysiological measures, in particular event-related potentials (ERPs), different paradigms have been created to impede conscious perception and thus investigate how visual awareness emerges. Such paradigms are designed either to interfere with the normal perception of stimuli (sensory (un)awareness), or to divert attention from them (attentional (un)awareness)^25^. The former paradigms include techniques such as stimulus masking^26^ (where stimuli are presented very briefly and are followed by a masker that prevents conscious detection), stimulus crowding^27,28^ (where a target is presented along with numerous other stimuli, causing it to go undetected), or interocular suppression^29^ (where different stimuli are presented simultaneously to each eye, allowing only one to be consciously detected). The latter include paradigms where attention is directed towards irrelevant aspects of the visual scene, away from the target information (e.g., inattentional blindness or attentional blink paradigms^30,31^; see Railo et al.^32^ for a review).

By comparing conscious and unconscious presentations of stimuli, different studies have attempted to identify the electrical brain responses associated with awareness. Two possible ERP markers have been put forward as possible electrical correlates of visual awareness. The first one to be highlighted is a late positive potential situated over centro-parietal electrode sites around 300-600 ms after the stimulus onset. This ERP wave, termed the P300, was found to emerge when participants detected stimuli in tasks manipulating visibility^33^. However, arguments against the P300 being the most reliable correlate of awareness have underlined that other factors, such as task relevance, may be critical for its appearance^34,35^. Moreover, the P300 has been found to correlate with processes separate from awareness, such as working memory^36,37^, context updating^38^, and post-perceptual processing^39^. Some studies have further shown that conscious report for a stimulus can take place before the onset of the P300^40^, shedding more doubt on the correlation between the P300 and awareness.

Subsequently, a number of observations have pointed to an earlier negative deflection over temporo-occipital regions that may in fact index awareness^41^. This earlier component is observed after ∼200 ms post-stimulus and presents as a greater negativity for conscious compared to unconscious stimulus presentations, consequently dubbed the Visual Awareness Negativity (VAN). The VAN has been found using different methods to manipulate awareness, even when controlling for the task relevance of the stimuli, objective task performance and a variety of types of attention (see Förster et al.^41^ for a review). This component has been posited to be the earliest correlate of visual awareness in the human brain^41,42,43^.

A controversy has emerged regarding which, if any, of these two markers actually reflects awareness, especially as they are loosely linked to two recent influential theories of awareness^44,45^. The global neuronal workspace theory^44^ postulates that awareness arises when sensory information, coded by modular cerebral networks, is amplified by attention and subsequently recruits neurons widely distributed in the brain. This process creates a “neuronal workspace” in which information becomes available for different complex processes, such as working memory, perceptual categorisation, or memorisation, allowing the emergence of awareness^44^. In this framework, awareness has been hypothesised to be indexed by the P300.

An alternate theory highlights the role of recurrent processing in enabling awareness^45^. This theory states that the feedforward sweep projecting information bottom-up through the cortical hierarchy is not sufficient to produce awareness. Rather, it is the subsequent feedback projections from higher to lower-tier areas following the initial feedforward sweep that are required to produce conscious perception^45^. Such feedback activity is largely localised to early regions receiving sensory (e.g., visual) information and suggests an earlier time scale for the emergence of awareness more akin to the VAN^41^.

Both theories have empirical support for their interpretations^41,46,47,48^ and the question remains open as to which one better accounts for perceptual awareness.

The discrepancies in findings regarding the electrical correlates of awareness and unconscious emotion processing may be due to the manner in which awareness is manipulated in previous paradigms. Indeed, awareness is generally impeded by modifying viewing procedures that, as noted above, are rarely if ever found under normal viewing conditions.

One way to circumvent this issue would be to allow the participants to explore the visual scene freely during the electrophysiological recordings. To our knowledge, unconstrained visual search has never been used in tasks measuring electrophysiological outcomes. Since natural viewing conditions entail saccades and multiple fixations that do not systematically produce awareness, we decided to take advantage of the so-called “normal blindness” or Look-But-Failed-To-See phenomenon^8^ to investigate the neural correlates of visual awareness and simultaneously to explore unconscious processing of emotional faces.

In the current study, participants were allowed to explore a visual scene freely, in search of a target stimulus. These visual scenes were composed of multiple faces displaying neutral, happy, and fearful expressions. Participants were asked to localise a single target face composed either of a fearful or a neutral expression in each trial (see Fig. 1a). Eye-movements and electroencephalography were recorded simultaneously during the search. The electrical potentials triggered at each fixation on the target face stimulus (the Fixation-Related Potentials or FRPs) were computed. Separate FRPs were obtained when the target was fixated but not reported (unaware condition), and when the target was fixated and subsequently reported (aware condition). These were further separated according to target expression (fearful vs. neutral).

**Fig. 1.**
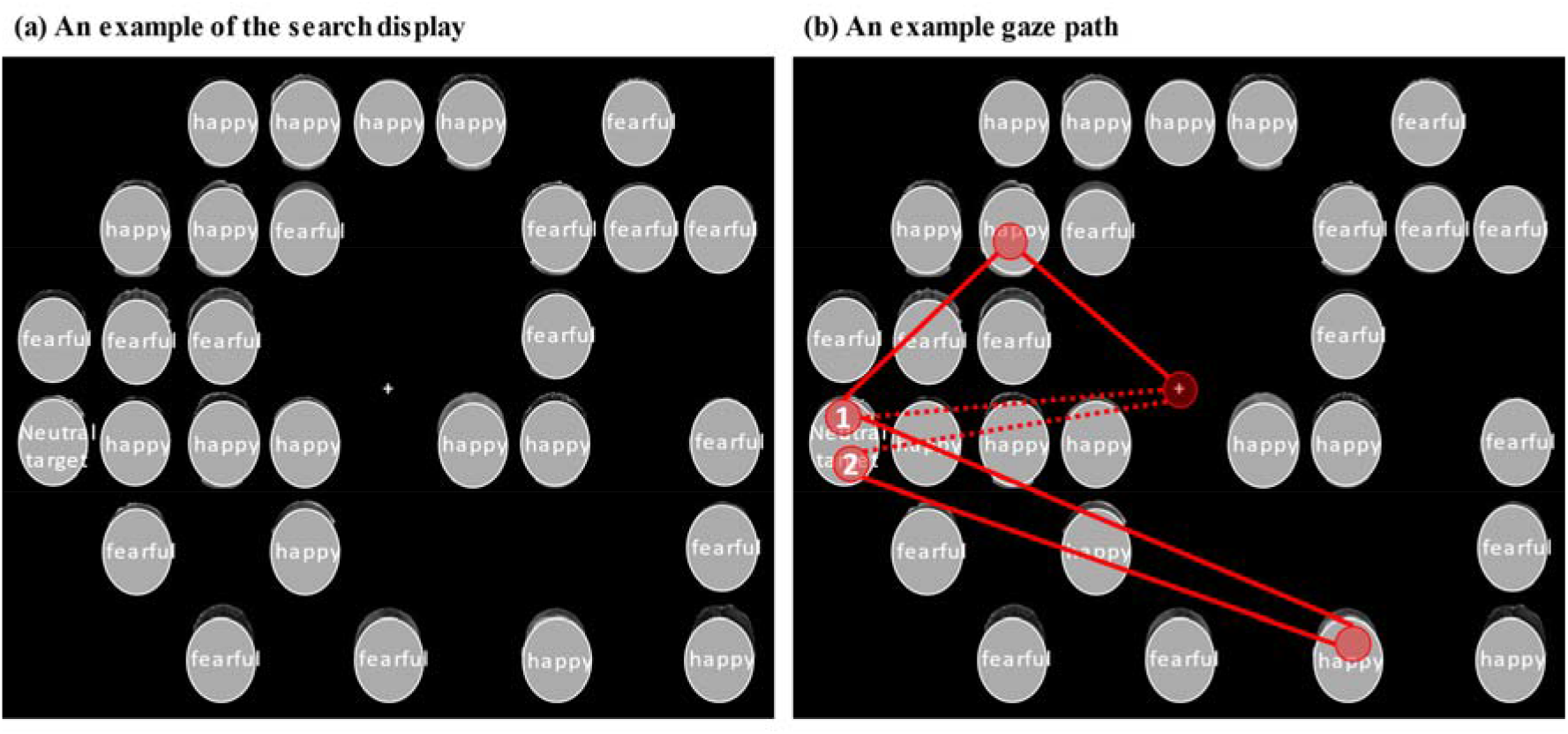
(a) An example of the search display in each trial. Here the target is a single neutral face amongst happy and fearful distractor faces. (b) An example gaze path. Only fixations on the targets are marked and analysed in this study. The number inside the circle indicates the order of the fixations on the same target in one trial. In this example, one or two fixations can be made on the neutral target face. If the first fixation on the target is followed by a saccade to another face image and is not followed by a report for the target, this fixation would be identified as an *unaware fixation*. Then, in this trial, the last fixation on the target followed by a saccade to the central fixation point (dashed line) and subsequently a correct target localisation would be identified as the *aware fixation*. If, however, the first fixation on the target is followed by a saccade to the central fixation point and a correct target localisation, it would be identified as the *aware fixation*, and there would be no unaware fixation from this trial. See the Method for a detailed description of the procedure. Note: The face stimuli are covered and de-identified in this picture in accordance with bioRxiv policies. Actual face images were used in the experiment.

On a trial, a variable number of unaware fixations could occur for fearful and neutral targets. The first such fixation on the target was used to compute the “unaware” FRP. The last fixation on the target comprised the “aware” FRPs, which necessarily terminated the search (see Fig. 1b). However, to ensure that this target fixation was not the last saccade and that it was followed by another saccade (as was the case for unaware fixations), participants were instructed to gaze at a specific location in the middle of the screen as soon as they saw the target, thereby activating the mouse cursor which allowed them to provide their manual response by clicking at the target location.

The aim of the current study was to identify the neural activity underlying conscious and, if present, unconscious processing of fearful and neutral faces, using free visual search for the first time. Moreover, regarding the neural correlate of awareness, we aimed to establish whether awareness arises early (∼200 ms) or later (>300 ms) in the stream of visual processing.

## Results

### Behavioural results: Accuracy and dwell time

The overall accuracy (proportion of correct responses) was 0.97 (*SD* = 0.03) at the target face localisation task. Due to the nature of the experiment, i.e., unlimited time to find a target, we expected a very high level of accuracy for all participants. No further analysis was performed on the accuracy data.

A 2(awareness: aware, unaware) x 2 (target emotion: fearful, neutral) repeated-measures ANOVA was performed on the average dwell time on targets. As shown in Fig. 2, a main effect of awareness was found, *F*(1,31) = 201.72, *p* < .001, *η*_*p*_^*2*^= 0.87, which was modulated by a significant interaction with target emotion, *F*(1,31) = 16.29, *p* < .001, *η*_*p*_^*2*^=0.34. Follow-up comparisons showed that, on unaware fixations, no difference was found between the dwell time on fearful targets and on neutral targets, *p* = .110. However, when participants were aware of the stimuli, dwell time on fearful targets was significantly shorter than neutral targets, *p* < .001.

**Fig. 2.**
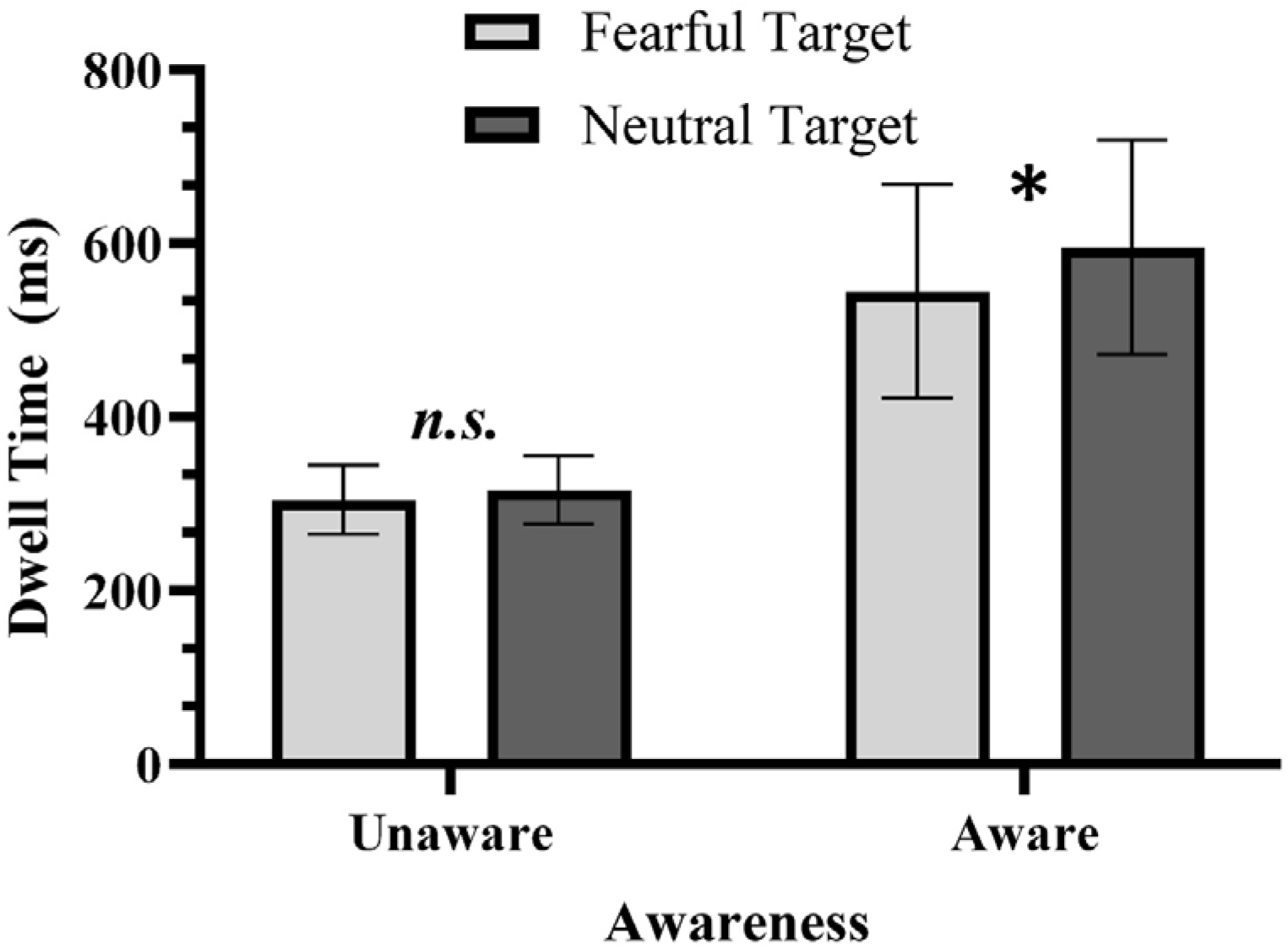
Dwell time data in different awareness conditions for fearful and neutral targets. Note: * *p* < .001.

### FRPs

The first Factorial Mass Univariate Analysis (FMUA) omnibus 2(awareness: aware, unaware) x 2(target emotion: fearful, neutral) ANOVA was performed on signals from posterior electrodes (P7/8, PO3/4, PO7/8, PO9/10, O1/2, Oz; for the FRP waveforms and topographic maps see Fig. 3a) to examine posterior components.

**Fig. 3.**
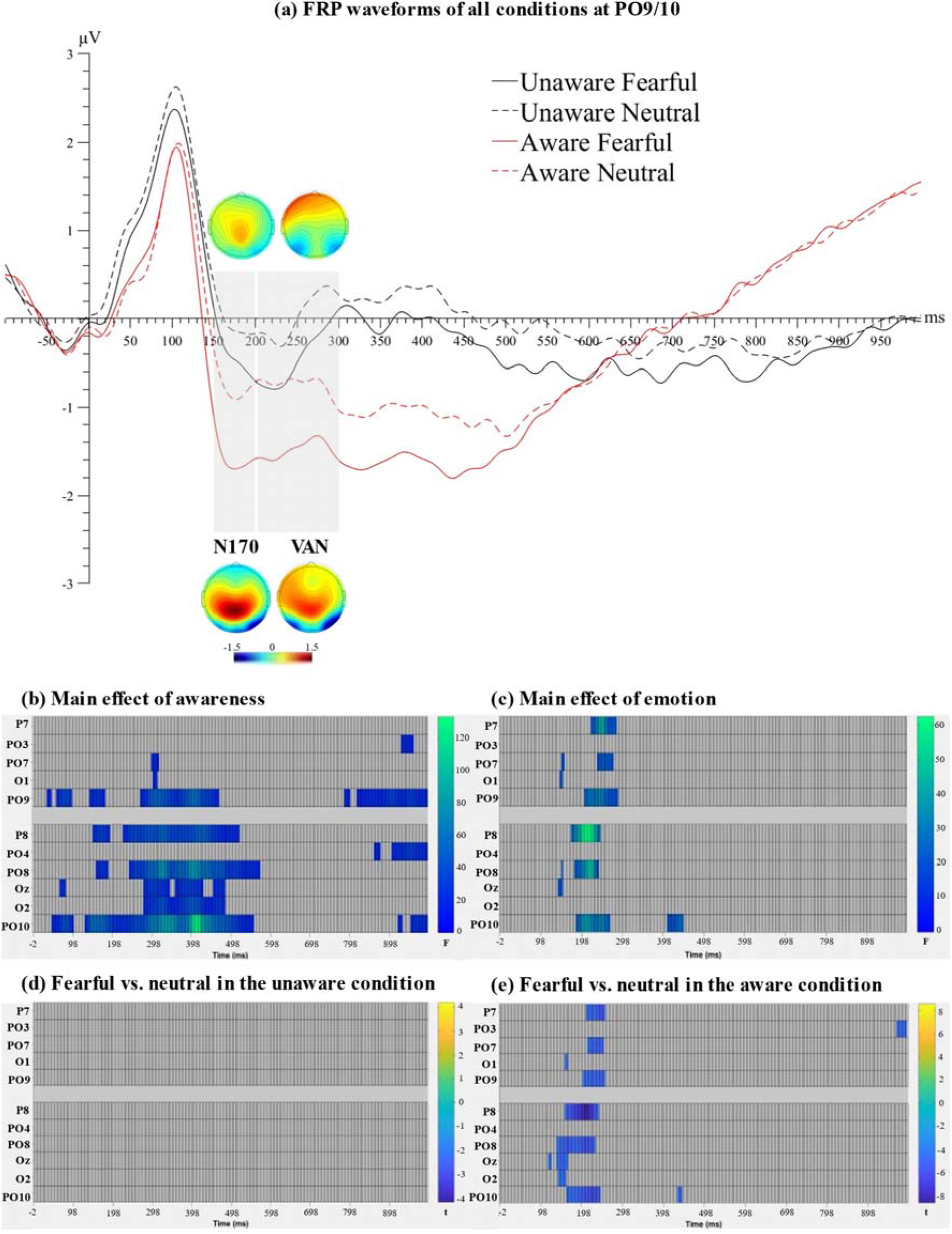
(a) FRP waveforms of all conditions at electrode PO9/10, the electrode sites where the maximal omnibus effect of awareness was found. Topographic maps for the highlighted time windows (the N170 and the VAN) were plotted separately for the unaware (top) and aware (bottom) conditions, collapsed across the emotion of the target face. Raster plots of FMUAANOVA results show (b) the significant main effect of awareness, (c) the significant main effect of emotion and (d & e) effects of emotion in different awareness conditions.

### The N170 and the VAN

The main effect of awareness was found to be significant in a face-specific N170 (150-200 ms) and the VAN (200-300 ms) time windows across multiple electrodes. Specifically, FRP amplitudes in the aware condition were more negative than the unaware condition at P8, PO8 and PO9/10 in a common time window spanning from 130 to 190 ms. Similarly, after 200 ms, signals in the aware condition were also more negative than the unaware condition, over most of the posterior electrodes (P8, PO7/8, PO9/10, O1/2, Oz). This negativity towards consciously processed stimuli spanned a large time window (e.g., 242-570 ms for PO8, Fig. 3b), which is not usually the case with a VAN that is often observed between 200-300 ms and peaks around 250 ms post-stimulus^41^. In the aware condition, participants had to maintain the location of the target face in their working memory before making a response. Specifically, they made their response after they had first fixated back at a central fixation and activated the mouse cursor (see Method). Therefore, differences between aware and unaware conditions not only encompassed effects of awareness but also likely effects of working memory, which oftentimes are reflected as later components (e.g., contralateral delay activity/CDA^36,37,49^). Based on previous literature on the VAN^41,42,43,50^, significant negativity towards aware stimuli as compared to unaware stimuli between 200-300 ms likely reflected the VAN in the current paradigm.

Additionally, the omnibus ANOVA revealed a main effect of emotion over 142-154 ms at PO7/8, O1 and Oz, and over 174-282 ms at P7/8, PO7/8 and PO9/10 (Fig. 3c), such that a fearful target was associated with more negative FRP amplitudes, compared to a neutral target, in time windows encompassing the N170 and the VAN. As part of our *a priori* contrasts, we compared the FRPs between fearful and neutral targets in the unaware and the aware conditions, respectively, using the Mass Univariate *t_max_* tests^51^. No emotion-related difference was found in the unaware condition (Fig. 3d). However, in the aware condition, a fearful target was associated with more negative amplitudes than a neutral target between 118-162 ms at PO8, O1/2 and Oz, and between 162-254 ms over P7/8, PO7/8 and PO9/10 (Fig. 3e). Therefore, the FRPs could distinguish a fearful from a neutral expression in the N170 and the VAN time range, only when participants were aware of the stimuli.

### The P300

A second FMUA omnibus 2(awareness: aware, unaware) x 2(target emotion: fearful, neutral) ANOVA was performed on parietal electrodes (Pz, P3/4, PO3/4) to examine any effect on the P300. We found a main effect of awareness with a significant positivity for aware compared to unaware stimuli between 300-590 ms on Pz and P3/4 (Fig. 4a & 4b). While such a parietal positivity towards aware stimuli may reflect an effect of awareness, it may also indicate working memory processes, as explained earlier. Supporting this, we found an unexpected late negativity from 738 ms onwards in these parietal regions in the aware condition (Fig. 4a & 4b), which aligns with the marker for information maintenance in working memory (CDA)^49,52^.

**Fig. 4.**
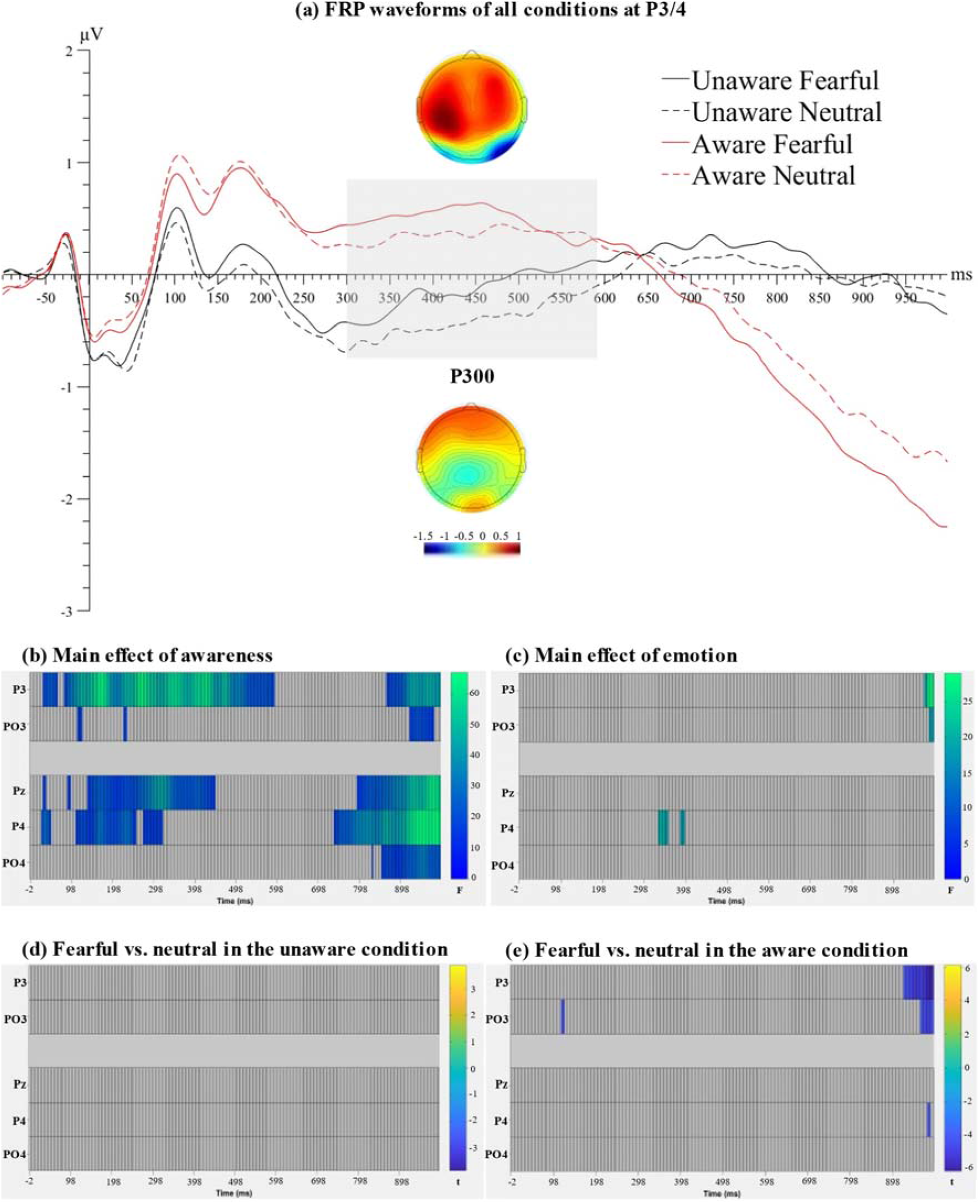
(a) FRP waveforms of all conditions at electrode P3/4, the electrode sites showing the maximal omnibus effect of awareness. Topographic maps for the highlighted time window (the P300) were plotted separately for the aware (top) and unaware (bottom) conditions, collapsed across the emotion of the target face. Raster plots of FMUA ANOVA results show (b) the significant main effect of awareness, (c) the significant main effect of emotion and (d & e) effects of emotion in different awareness conditions.

The omnibus ANOVA on parietal electrodes also revealed a main effect of emotion (Fig. 4c). We conducted *a priori* contrasts between fearful and neutral target conditions at each level of awareness. As shown in Fig. 4d and 4e, no emotion-related difference was found in the P300 time window in either awareness condition. However, towards the end of the epoch (926-994 ms over P3 and 966-994 ms over PO3), a fearful target face was associated with more negative amplitudes than a neutral target face in the aware condition, likely reflecting an enhanced working memory consolidation for fearful target faces relative to neutral ones.

### Conscious report occurs with gradual increase in the P300/CPP

In most trials, there are multiple or repeated fixations on the target. Thus, the current paradigm provides an opportunity to investigate the electrical activity across the repeated fixations prior to the final conscious report for the target. To test whether perceptual evidence was accumulated in a gradual or an all-or-none manner leading to a conscious report, we extracted trials where participants reported seeing the target face after at least three fixations on the target. Then, we compared the FRPs between the last three fixations on the target (i.e., the last fixation, 1-prior, 2-prior). In keeping with previous literature on evidence accumulation^53,54^, we selected the centro-parietal electrodes as regions-of-interest for the known electrophysiological evidence accumulation signal, the central-parietal positivity (CPP)^54^. A one-way (fixation: the last fixation, 1-prior, 2-prior) FMUA ANOVA was performed on Cz, Pz, P3 and P4. We found a main effect of awareness over all four electrodes (see Fig. 5a & 5b). Follow-up *t*-tests showed a significant positivity for the last fixation, compared to both 1-prior (154-282 ms; Fig. 5c) and 2-prior fixations (154-302 ms and 342-362 ms; Fig. 5d). Additionally, FRPs for the 1-prior fixation were significantly more positive than for the 2-prior fixation, between 398-430 ms at Cz (Fig. 5e). Therefore, the CPP seemed to increase gradually from 2-prior to the last fixation that preceded a conscious report for the target.

**Fig. 5.**
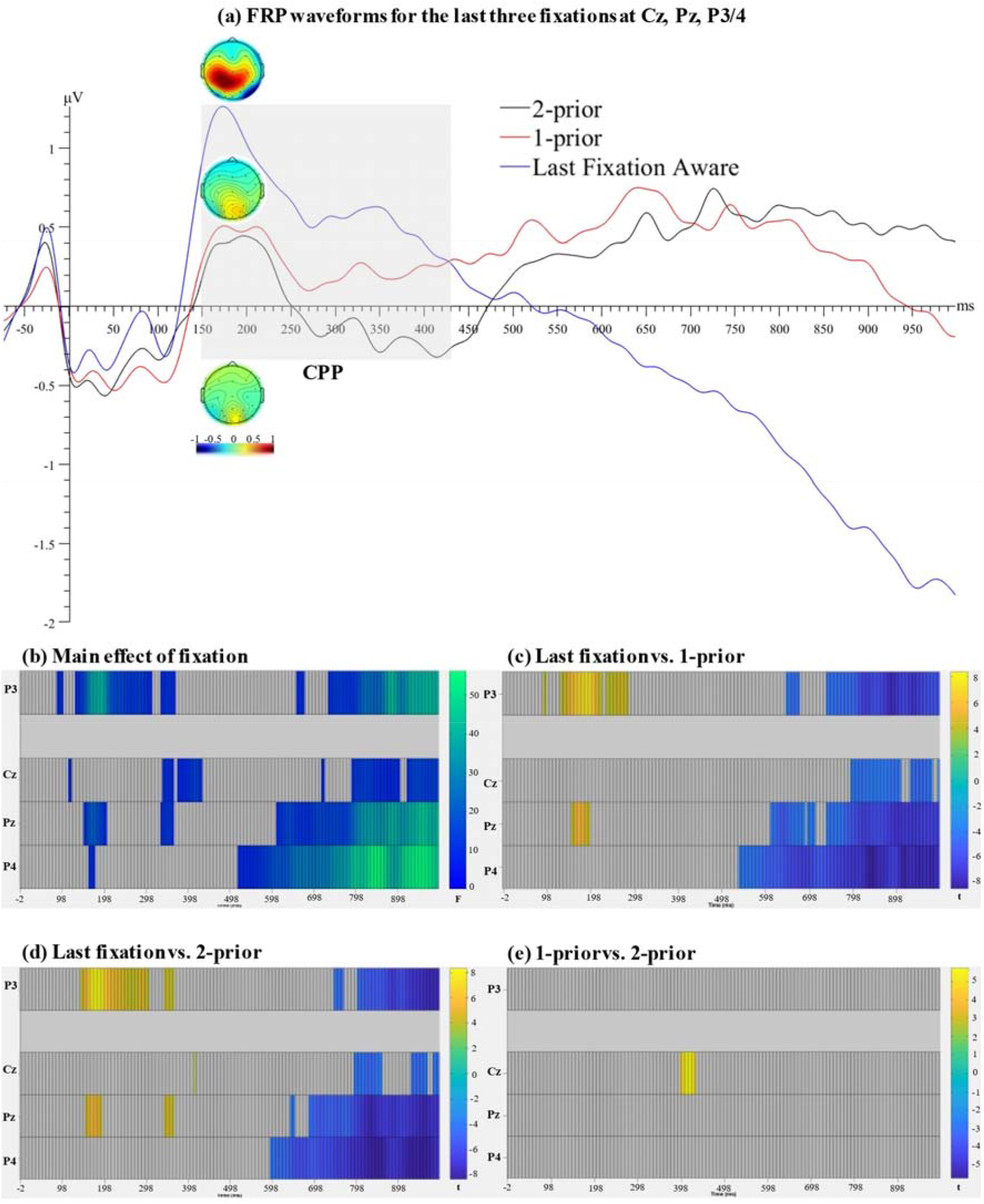
(a) FRP waveforms for the last three fixations at Cz, Pz, P3/4. Topographic maps for the highlighted time window (150-430 ms) were plotted separately for different fixation conditions. Raster plots of FMUA ANOVA results show (b) the significant main effect of fixation, and (c-e) follow-up pairwise comparisons of the fixation effects.

## Discussion

This study examines awareness and unconscious processing of emotional faces under free-viewing conditions, using combined EEG and eye-tracking methodologies. Effects of awareness were found at several time windows including the N170, VAN and the P300, while effects of emotion (in the aware condition) were found mainly in the N170 and VAN time windows. No differences were found between expressions in the unaware condition. Furthermore, the differences in the neural activity between aware and unaware conditions were consistent with fixation dwell time. Specifically, participants fixated at the target face for a longer time when they were aware of, and subsequently reported the target, compared to when they did not report it.

In the current paradigm, visual information was identical across aware and unaware conditions as the visual scene remained unchanged throughout each visual search trial. Indeed, visual scenes during unaware fixations on the target were identical to those that led to a response, such that the two conditions differed only in the fact that they entered consciousness in the latter case. Still, one could contend that targets were insufficiently differentiable from the distractors. Viewers would thus have processed the stimulus consciously but pursued their search due to their uncertainty. However, in a number of trials, participants correctly localised the target face upon the first fixation on the target, and the task was judged as relatively easy with participants showing high accuracies on the task (97%), which argue against this possibility.

Compared to the unaware fixations, those leading to conscious report produced more negative FRPs starting at ∼130 ms. Previous observations have provided that visual awareness produces a negative ERP deflection over temporo-occipital electrodes, which emerges sometime after 100 ms and peaks at ∼250 ms^41^, when comparing aware and unaware stimulus presentations. This deflection has been observed using different methods to interfere with conscious processing, including masking, stimulus degradation, attentional blink, change blindness and bistable perception (see Koivisto and Revonsuo^55^ for a review).

However, to date, no attempt had ever been made to determine if this is applicable to conditions of free visual search. Here, the evidence indicates that such a negativity does indeed index awareness, and further demonstrates that the effect is longer-lasting than in constrained viewing conditions^41^.

At later time points (beyond ∼300 ms), our results show a greater positivity over the parietal vertex region for aware stimuli, corresponding to the P300. As noted above, the P300 has also been suggested to be a marker of awareness as it was initially reported when comparing conscious and unconscious processes^33,47^. It has been proposed that once a certain threshold is reached, numerous cortical regions are recruited and the sensory information becomes accessible by different information processing modules, leading to awareness^33^. This view has been supported notably by the observation that cortical activation during the P300 encompasses extensive parietal and frontal brain regions.

However, different from the earlier negativity, the P300 has been found to be linked to post-perceptual processes subsequent to awareness. Some have therefore argued against the candidacy of the P300 for awareness. For example, using a masking procedure in an attentional task, Del Zotto and Pegna^56^ observed a negativity in the 200 ms range for consciously detected emotional faces. Interestingly, the P300 was modulated depending on whether the face was a target or not, revealing that the magnitude of this marker depends on stimulus processes that are separate from awareness. Elsewhere, Pitts and colleagues compared the ERPs to geometric patterns that participants were or weren’t aware of, using an inattentional blindness task^57^. Compared with the unaware condition, the aware condition showed an increased negativity around 200-240 ms, however the P300 appeared only when the geometric shapes were relevant to the task at hand^57^. These results link the P300 to the relevance of the stimulus as a target rather than awareness *per se*.

Building on this idea, the P300 has been posited to be linked to template-matching in search tasks. Its amplitude would indicate how well a stimulus is recognised and identified as a target^58,59,60^. When participants are asked to detect a target, an internal template must be maintained for comparison with the stimulus under scrutiny. The greater the similarity between the target and the template, the larger the amplitude of the P300^61^. In line with these views, the P300 here could be seen as the neural activity indexing the goodness-of-fit between with the internally held template and the external stimulus. In this interpretation, sensory evidence is accumulated until subjective perceptual evidence reaches a threshold where a decision can be made^54^. Our observations validate this view: when comparing the last fixation that led to awareness with the previous fixation and the one before that, the P300, or more precisely the CPP, is seen to increase gradually. This finding is consistent with the idea of accumulating evidence of a match between the template and the fixated stimulus.

One additional function is also likely indexed by the P300 in this study: working memory. The current task required explicit target identification, as well as the encoding of the spatial location of the target in working memory, as target location had to be subsequently indicated using the mouse. The P300 has been found to increase with working memory load^36,37,49^, suggesting that the manner in which task responses were provided contributed partly to the enhancement of this component. Along these lines, we also observed a late negativity at parietal regions in the aware, compared to the unaware condition, likely reflect information maintenance in working memory^36,37,49^.

To summarise, during visual search, an early negativity emerging at ∼130 ms is likely the first electrical correlate for awareness of target information whilst the P300 or CPP appears to reflect evidence accumulation across target fixations, possibly through an ongoing comparison/matching between the stimulus and a target template.

Furthermore, differences were found between the FRP responses to different target expressions. Specifically, fearful target faces were associated with a greater negativity over the N170 and the VAN time windows compared to neutral target faces. Critically, the FRPs were increased by target fearful expressions only when participants were aware of the stimuli.

The enhanced negativities in the aware condition over the 170 and 200 ms time periods for fearful faces are unsurprising and are consistent with previous observations reporting an N170 enhancement and an early posterior negativity (EPN) for emotional expressions^42,62,63,64^ (see Schindler & Bublatzky^65^ for a review). However, the results found in the current visual search task indicate that no apparent processing of fearful expressions arises when stimuli are fixated but not consciously detected.

Nevertheless, with no electrical difference observed for unaware emotions, the absence of any unconscious processing cannot be unequivocally asserted. Indeed, the build-up of the P300/CPP with repeated target fixations demonstrates that some unconscious processing of the target faces occurs during visual search, although the findings do not suggest any differentiation of this activity across expressions.

Finally, the dwell time on a fearful target face was shorter than that on a neutral target face in the aware condition, while this was not observed in the unaware condition. The shorter dwell time on fearful face targets could suggest a more rapid processing of fearful expressions, compared to neutral ones, possibly due an easier detection of fear than of neutrality. Alternately, when searching for a fearful face, neutral face distractors may have constituted less of a hindrance (i.e., did not capture attention) than fearful face distractors when searching for the neutral face target. Evidence for both these possibilities has been found^12,66,67,68^, and both are likely to have affected the visual search dwell times, possibly also in conjunction with one another.

In conclusion, the evidence obtained in this study suggests that during free viewing visual search, some non-specific processing of faces arises without awareness. When the sensory system has accumulated sufficient evidence regarding the emotional expression (indexed by the P300/CPP), awareness emerges at around 130 ms, with a specific and conscious processing of emotions emerging in the N170 time range and beyond.

## Method

### Participants

The sample size was determined based on the smallest effect size reported in our previous work on awareness of fearful faces (*η*_*p*_^*2*^= 0.33)^50^. For our within-participants 2-by-2 factorial design, the minimum sample size was 16 for a significant main effect of awareness that is sufficiently powered (i.e., 90%), with an effect size of 0.33 at an alpha level of 0.05, two-tailed (calculated with MorePower Software^69^). Out of an abundance of caution, especially considering that this is a novel paradigm, we doubled the sample size. Thirty-two participants (*M_age_* = 21.4 years, *SD_age_* = 3.1 years; 12 males, 20 females) were recruited at the University of Queensland and were compensated with either course credits or $50 (AUD) for their participation. All participants had normal or corrected-to-normal vision and they had no history of neurological or psychiatric conditions. After data pre-processing, data from two participants was excluded for the FRP analyses (see Data pre-processing). As a result, the final sample for the FRP analyses consisted of 30 participants (*M_age_* = 21.2 years, *SD_age_* = 2.9 years; 11 males, 19 females). The experimental procedure was approved by the ethics committee of the University of Queensland. All participants provided informed consent prior to their participation.

### Apparatus and stimuli

All stimuli were presented on a 19” colour LCD monitor (resolution: 1280 × 1024 pixels) with a viewing distance of 65 cm. The experiment was programmed and run in PsychoPy3^70^.

We obtained fearful, happy and neutral face images from 48 models (24 males, 24 females) from the Radboud Faces Database^71^. Each face image was rendered black-and-white and scaled approximately 4.2 cm x 3 cm (3.7° x 2.6° in visual angle; see Fig. 1a). The average luminance was matched across images. All photo editing was done in Photoshop 2021 version 22.4.0 (Adobe Systems, San Jose, CA).

Each search array consisted of one target face and 28 distractor faces. The screen was divided into 54 (a 9-by-6 grid) virtual image holders. In each search array, images were presented at 29 image holders randomly selected by the experimental program. To make sure that the task was sufficiently difficult, we used two non-target expressions in the distractor face images (Fig. 1a). Specifically, in the fearful target face block, a target fearful face was presented among 14 happy distractor faces and 14 neutral distractor faces. In the neutral target face block, a target neutral face was presented among 14 happy distractor faces and 14 fearful distractor faces.

### Eye movement and EEG data acquisition

Monocular gaze position was recorded using the Eyelink 1000 plus system (SR Research Ltd., Canada) with a spatial resolution of <0.01° and a sampling rate of 1000 Hz.

Continuous EEG was acquired at 500 Hz using the BrainProducts 32-channel system (Brain Products, Germany) using the international 10–20 configuration. During recording, EEG signals were band-pass filtered between 0.01-40 Hz, and a notch filter of 50 Hz was used to reject power line noise. Recordings were referenced online to a reference electrode taped to participants’ left ear. Impedances were kept below 15kΩ

#### Procedure

Before the experiment began, the eye tracker was calibrated with a 9-point calibration, and participants completed five to ten practice trials with one target expression (fearful or neutral), randomly determined by the program.

As shown in Fig. 6, each trial started with a Ready screen where participants were presented with a text prompt (“Press SPACE bar when you’re ready.”). Once participants pressed the Space bar, a fixation screen was presented. Participants were required to stably fixate at the central fixation cross for 500 ms to proceed. Afterwards, participants were presented with a search array and were asked to move their eyes across the screen to find the target face as quickly as possible. Once they found the target, they were required to fixate back at the central fixation cross and press the Space bar to activate the mouse cursor. Then, they could move the cursor to the target face and click on it to indicate that they had found it. The purpose of requiring participants to fixate back at the central fixation prior to making a response (i.e., clicking on the target) was to prevent any motor preparations or movements from contaminating the FRP data in the time window of interest (i.e., 0-1000 ms after the fixation on the target face) and to ensure that target detection was always followed by a saccade, as with an unaware/unreported fixation. After participants made their response, the search array was replaced by a blank screen of 1000 ms, which ended the trial.

**Fig. 6.**
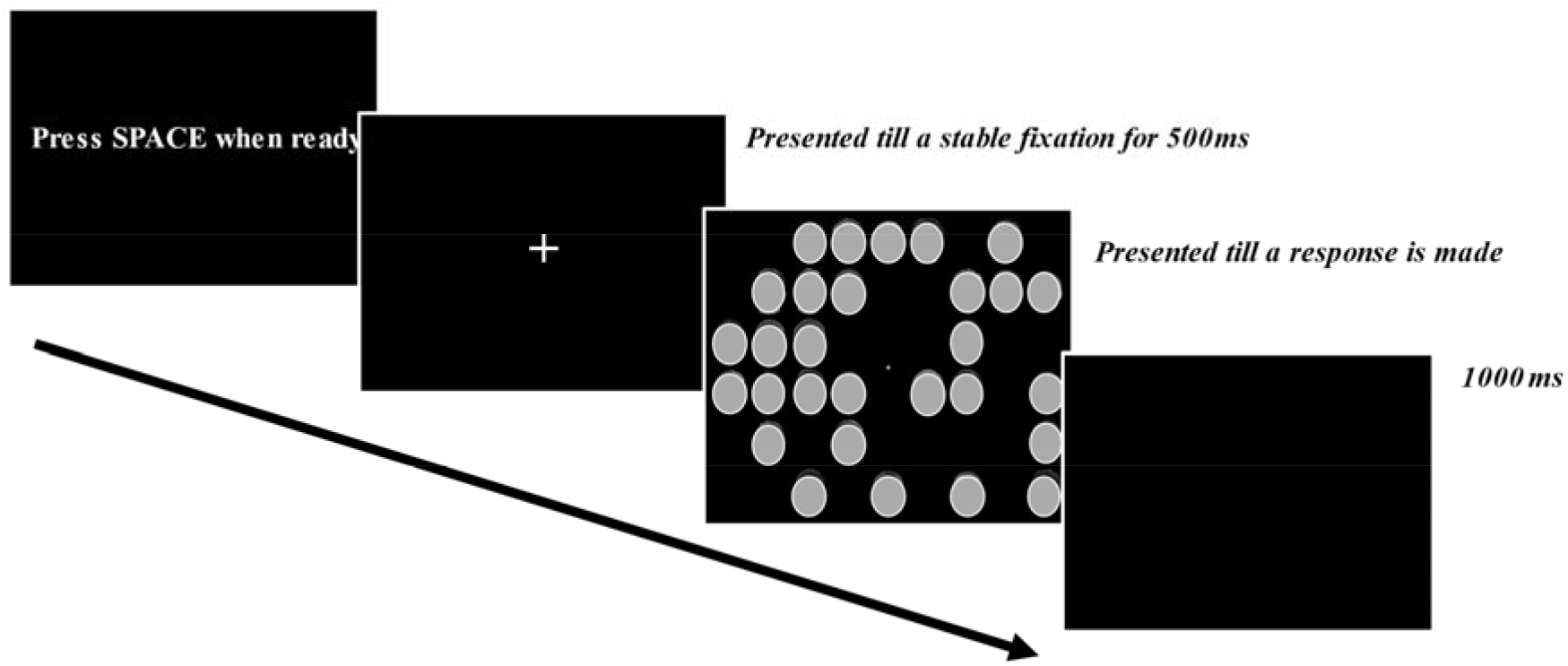
Time-course of events during a trial.

Participants performed two fearful-face-target blocks and two neutral-face-target blocks with 100 trials in each block. Breaks were allowed every 50 trials. The order of fearful and neutral blocks was randomised, separately for the first and the second half of the experiment.

### Data pre-processing

Pre-processing of the EEG data was performed with EEGLAB^72^ and ERPLAB^73^. We interpolated individual electrodes that produced sustained noise signals throughout the experiment. Signals were filtered from 0.1 to 30 Hz, and a notch filter of 50 Hz was included to remove line noise. We re-referenced the signals to the average of all electrodes. FRP signals were segmented into epochs with a time window of 1000 ms from the onset of a fixation on the target face, relative to a pre-fixation baseline (−100 to 0 ms). A fixation was identified as within the target face region if the distance between the fixation and the centre of the image was smaller than half of the diagonal of the image holder (95 pixels).

From each trial, one or two fixations on the target could be analysed (see Fig. 1b). Specifically, a fixation on the target was identified as an aware fixation if it was immediately followed by a saccade to the central fixation point and subsequently a correct localisation of the target. The epoch time-locked to this fixation is labelled the aware condition. A fixation was identified as an unaware fixation if it was the first fixation on the target in each trial but was not followed by a saccade to the central fixation point in preparation for making a response. Both conditions were separated by the target face expression (i.e., fearful vs. neutral).

In order to correct for eye-movements or eye-blinks components in the EEG data, we ran an optimised Independent Component Analysis (ICA) guided by the eye-tracking data, using the EYE-EEG toolbox^74^ and customised MATLAB scripts. Here briefly, we first parsed the eye-tracking data and re-sampled it from 1000 Hz to 500 Hz to match the EEG data sampling rate. Then, we synchronised the eye-tracking and EEG data by identifying and matching the onset of the triggers in both data. Bad eye-tracking data including intervals of eye-blinks and out-of-range eye-movements were detected and marked. Afterwards, saccades and fixations were detected using the velocity-based saccade detection algorithm^74,75^. Specifically, (micro)saccades were defined as intervals in which the velocity of the recorded eye movements exceeded six median-based standard deviations of all eye velocities for at least four samples. Additionally, for micro-saccades detection, a magnitude threshold of 1° was used, and the interval between successive saccades was set as 50 ms for saccades clustering^74,76^.

A customised ICA was then run on the EEG data that contained eye-movement information, using the infomax ICA algorithm of Bell and Sejnowski^77^. A detailed procedure has been described elsewhere^78^. After eye-movement and eye-blink components were decomposed and removed from the EEG data, we segmented the data based on the fixation events (described above). A threshold of −80 to 80 µV was used for automatic detection of artefacts in the segmented data. We further inspected the data and rejected epochs containing artefacts on a trial-by-trial basis. Epochs containing bad eye-tracking intervals were automatically removed. As it was important for us to remove eye-related components from the EEG data, we excluded participants with eye-related components (eye-blinks and eye-movements) identified with a likelihood of less than 50%. As a result, data from two participants were excluded from further FRP analyses.

The remaining participants had on average the following mean numbers of epochs per condition: 87 epochs (*SD* = 18) for unaware fearful targets, 143 epochs (*SD* = 22) for aware fearful targets, 101 epochs (*SD* = 19) for unaware neutral targets, and 138 epochs (*SD* = 23) for aware neutral targets.

We also obtained target face dwell time data for all conditions. Specifically, dwell time was calculated as the time between the onset of the first saccade into the target face region to the onset of the saccade leaving the target face region. Dwell times shorter than 50 ms or longer than 1500 ms were excluded.

All codes used in our data pre-processing and analyses can be found at https://osf.io/9hswj.

### FRP data analysis

The FRP data were analysed under the Factorial Mass Univariate Analysis (FMUA) framework^79^. For our main analyses, separate FMUA were conducted over all time-points in the FRP epoch (i.e., 0-1000 ms) on posterior electrode sites (P7, P8, PO3, PO4, PO7, PO8, PO9. PO10, O1, O2, Oz) encompassing the N170 and the VAN, and parietal electrode sites (Pz, P3, P4, PO3, PO4) encompassing the P300. Multiple comparisons were corrected using the *F_max_* approach, which estimates a null distribution of the maximal effect (i.e., the maximal *F* value) across the selected electrodes and time-points using a permutation test (10,000 permutations)^79,80^. The observed *F* statistic at each time point was considered significant if its value exceeded the 95th percentile of the null distribution. Pairwise follow-up comparisons were conducted using *t_max_* tests under the Mass Univariate Analysis framework using 10,000 permutations, with a two-tailed family-wise α level of 0.05^51^.

We additionally ran an exploratory analysis over all electrodes and all time-points in the epoch using the cluster-based permutation test, which offers better power for exploratory analyses^79^. The full report of the exploratory analysis is provided in the SupplementaryMaterials.

