## Supplementary Materials for "Now you see it: fixation-related electrical potentials during a free visual search task reveal the timing of visual awareness"

**Supplementary Results: A whole-brain exploratory analysis**

We performed a whole-brain exploratory factorial mass analysis on all time-points and all electrodes using the cluster-based permutation test to control for multiple comparisons (10,000 permutations). Electrodes were considered as spatial neighbours if they were within approximately 5.3 cm of one another, resulting in each electrode with on average 3.4 spatial neighbours. The cluster formation threshold was set at 0.05 and significant differences were determined with an alpha level of 0.05.

The FMUA omnibus 2(awareness: aware, unaware) x 2(target emotion: fearful, neutral) ANOVA revealed a main effect of awareness on all electrodes except FPz and Cz, extending throughout the whole epoch (cluster *p* < .001). The effect was the strongest over parietal and occipital sites (spatial peak at PO10) with a temporal peak at 410 ms, see Figure S1a.

The main effect of emotion was also significant on almost all electrodes (see Figure S1b), extending from 106 to 578 ms, with the effect being the strongest over occipital and parietal sites (cluster *p* = .001, spatial peak at P8). The temporal peak for the cluster was 210 ms.

The interaction between awareness and target emotion was non-significant, see Figure S1c. We conducted *a priori* contrasts between fearful and neutral targets using cluster-based permutation *t*-tests for the aware and the unaware conditions, separately. As shown in Figure S1d, in the unaware condition, no effect of emotion was found. In the aware condition, however, fearful compared to neutral target faces were associated with more negative signals between 102 and 482 ms over parietal and occipital electrodes (cluster *p* = .008, spatial peak at P8) with a temporal peak at 210 ms. An additional frontal positive cluster was found for fearful targets, spanning from 98 to 274 ms with a temporal peak at 210 ms (cluster *p* = .047, spatial peak at C3), see Figure S1e.

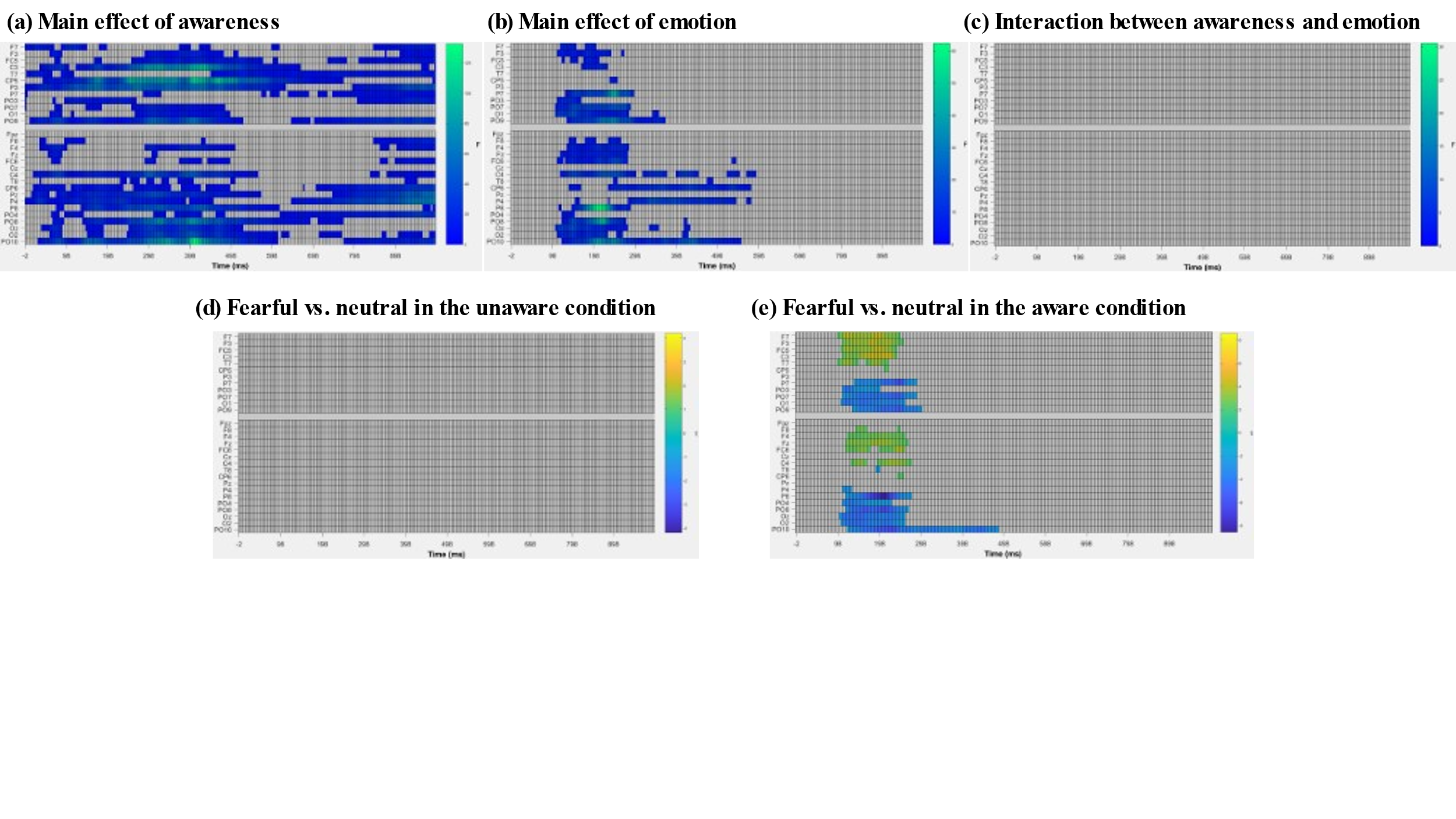
***Figure S1.*** Raster plots of FMUA ANOVA results show (a) the significant main effect of awareness, (b) the significant main effect of emotion and (c) non-significant interaction between awareness and emotion, and (d & e) raster plots for the fearful and neutral contrasts in different awareness conditions.
